# Dopamine receptors in obesity: a meta-analysis

**DOI:** 10.1101/2020.12.29.424782

**Authors:** Kyoungjune Pak, Myung Jun Lee, Keunyoung Kim, In Joo Kim

## Abstract

The brain plays a major role in controlling the desire to eat. This meta-analysis aimed to assess the association between dopamine receptor (DR) availability, measured using positron emission tomography, and obesity. We performed a systematic search of MEDLINE (from inception to June 2019) and EMBASE (from inception to June 2019) for articles published in English using the keywords “dopamine receptor,” “obesity,” and “neuroimaging.” Fitting a locally weighted estimated scatterplot smoothing curve for binding potential (BP_ND_) of the ventral striatum measured by ^11^C-raclopride was performed. Five studies involving 119 subjects were included in this analysis. ^11^C-raclopride was used in four studies, regardless of the ROIs included. The curve of BP_ND_ of the ventral striatum went up and down at BMI of 20–30 kg/m^2^ and became flat after a BMI of >40 kg/m^2^. Linear regression analysis was performed with BP_ND_ of the striatum from two studies (n = 20). The BP_ND_ of the striatum was negatively associated with BMI. In conclusion, dopamine plays a major role in the reward system with regard to obesity. Compared to normal-weight subjects, overweight and obese subjects had decreased DR availability. However, this tendency should be interpreted carefully with regard to radiopharmaceuticals and ROIs.

## INTRODUCTION

The prevalence of obesity has nearly tripled worldwide since 1975 (Organization, 2018). Obesity is a risk factor for several malignancies of the colon (Na & Myung, 2012), pancreas (Gukovsky, Li, Todoric, Gukovskaya, & Karin, 2013), thyroid (Mijovic, How, & Payne, 2011), liver (Alzahrani, Iseli, & Hebbard, 2014), and uterus (Gu, Chen, & Zhao, 2013) as well as for cardiovascular diseases (Burke et al., 2008). Obesity is caused by an imbalance between energy intake and expenditure over a long period of time (Morton, Meek, & Schwartz, 2014). This imbalance can be triggered by homeostatic, genetic, environmental, psychological, social, and cultural factors (Berridge, 1996; Pecina, Smith, & Berridge, 2006; Volkow, Wang, Tomasi, & Baler, 2013). There are phenomenological similarities between overeating in obesity and excessive drug use in addiction (Kenny, 2011); hence, the brain plays a major role in controlling the desire to eat (Pak, Kim, & Kim, 2018; Val-Laillet et al., 2015). Among the neurotransmitters synthesized and released at synapses in the brain, dopamine plays a major role in the executive function, motor control, and motivation and reward pathways, particularly in the regulation of food intake and body weight (Baik, 2013; Lam, Garfield, Marston, Shaw, & Heisler, 2010; Ravussin & Bogardus, 2000).

There is no direct method to measure dopamine levels in the human brain. Positron emission tomography (PET) and single-photon emission computed tomography using radiopharmaceuticals have been used to investigate the role of the dopamine system in obesity (Pak et al., 2018). Previous PET studies using dopamine receptor (DR) radiopharmaceuticals have shown varying DR availability (lower (Wang et al., 2001), higher (Gaiser et al., 2016), or not significantly different (Eisenstein et al., 2013)) in obese subjects compared to lean controls. Nevertheless, the exact role of DRs in obesity is unclear. In a previous study, van Galen et al. compared the DR availability in obese and lean subjects, regardless of radiopharmaceuticals (van Galen, Ter Horst, Booij, la Fleur, & Serlie, 2018). There was a non-linear relationship between the body mass index (BMI) and DR availability; the DR availability increased in moderate obesity (BMI: approximately 35 kg/m^2^) and the DR availability decreased with the progression of obesity (BMI: >40 kg/m^2^) (van Galen et al., 2018). Although the mean BMI and binding potential (BP_ND_) values are the best representatives of the association between obesity and DR availability, there might be a difference from the real interrelation between BMI and BP_ND_ in each subject. Therefore, we performed a meta-analysis to assess the association between DR availability, measured using PET, and obesity.

## METHODS

### Data Search and Study Selection

We performed a systematic search of MEDLINE (from inception to June 2019) and EMBASE (from inception to June 2019) for articles published in English using the keywords “dopamine receptor,” “obesity,” and “neuroimaging.” All searches were limited to human studies. The inclusion criteria were neuroimaging studies of DR availability in the striatum (the striatum, caudate nucleus, putamen, nucleus accumbens [NAc], and ventral striatum [VST]) in association with BMI. Reviews, abstracts, and editorial materials were excluded. Further, duplicate articles were excluded. If there was more than one study from the same institution, the study reporting information most relevant to the present study was included. Two authors performed the literature search and screened the articles independently, and discrepancies were resolved by consensus.

### Data Extraction and Statistical Analysis

Two reviewers independently extracted the following information from the reports: first author, year of publication, country, radiopharmaceuticals, number of subjects, BMI, and DR availability expressed in BP_ND_. In this meta-analysis, the association between BMI and BP_ND_ was investigated using two methods. First, BMI and the corresponding BP_ND_ were extracted from figures of each study using Engauge Digitizer, version 12.1 (http://digitizer.sourceforge.net), and plotted for both radiopharmaceuticals and regions of interest (ROIs). Linear regression analysis was performed to describe the relationship between BMI and BP_ND_. Fitting a locally weighted estimated scatterplot smoothing (LOWESS) curve for BP_ND_ of the VST measured by ^11^C-raclopride was performed. Second, we extracted the mean BMI and BP_ND_ values directly from each study, if provided by the authors, and plotted them for each radiopharmaceutical. If they were not provided by the authors, the mean BMI and BP_ND_ were calculated after extracting the BMI and corresponding BP_ND_ from the figures. Data from each study were analyzed and plotted using Prism 8 for macOS, version 8.3.1. (GraphPad Software, LLC, San Diego, CA, USA).

## RESULTS

### Study Characteristics

The electronic search identified 389 articles. Conference abstracts, animal studies, non-English studies, and studies that did not meet the inclusion criteria after screening the title or abstract were excluded. After reviewing the full text of 43 articles, nine studies including 205 subjects were eligible for inclusion in this study (Caravaggio et al., 2015; Dunn et al., 2010; Gaiser et al., 2016; Guo, Simmons, Herscovitch, Martin, & Hall, 2014; Karlsson et al., 2015; Karlsson et al., 2016; Steele et al., 2010; Volkow et al., 2008; Wang et al., 2001). Five studies using ^11^C-raclopride (Karlsson et al., 2015; Karlsson et al., 2016; Steele et al., 2010; Volkow et al., 2008; Wang et al., 2001), one using ^11^C-PNHO (Gaiser et al., 2016), two using ^18^F-fallypride (Dunn et al., 2010; Guo et al., 2014), and one using both ^11^C-raclopride and ^11^C-PNHO (Caravaggio et al., 2015) were included. The BP_ND_ of the VST, caudate nucleus, and putamen were extracted from four studies (Dunn et al., 2010; Gaiser et al., 2016; Karlsson et al., 2015; Karlsson et al., 2016), and the BP_ND_ of the NAc, caudate nucleus, and putamen were extracted from one study (Guo et al., 2014). NAc is the main component of VST (Heimer, 1978); hence, we included the BP_ND_ of NAc while analyzing the BP_ND_ of VST. Three studies included the preoperative and postoperative results of patients who had undergone bariatric surgery (Dunn et al., 2010; Karlsson et al., 2016; Steele et al., 2010). Figure 1 shows the detailed procedure. Table 1 summarizes the study characteristics.

**Figure 1.**
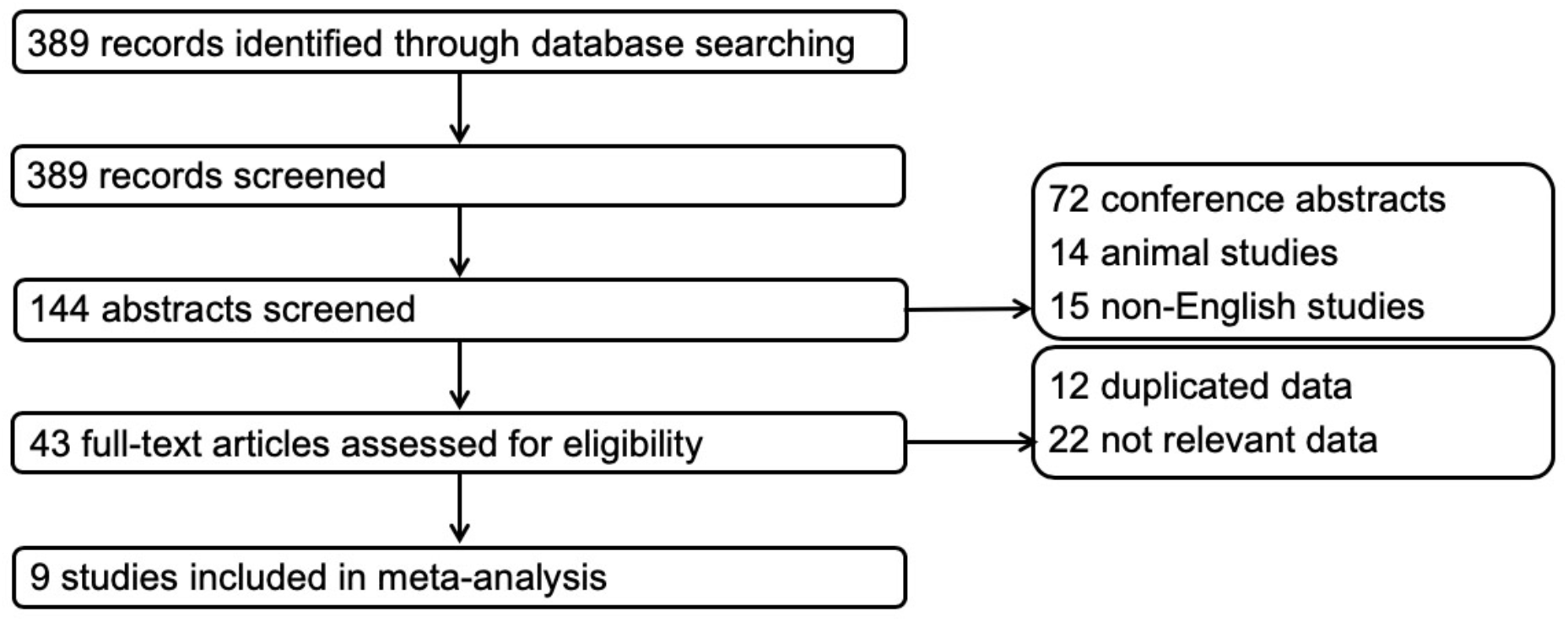
Flowchart.

**Table 1.** Studies included in the meta-analysis.

| Author | Year of publication | Country | Radiopharmaceutical | No. of subjects | Analysis 1 |  | Analysis 2 |  |
| --- | --- | --- | --- | --- | --- | --- | --- | --- |
|  |  |  |  |  | Source | ROI | Source | ROI |
| Caravaggio | 2015 | Canada | <sup>11</sup>C-PHNO | 26 | Fig 1 | VST | Fig 1 | VST |
|  |  |  | <sup>11</sup>C-Raclopride | 35 | Fig 3 | VST | Fig 3 | VST |
| Karlsson | 2015 | Finland | <sup>11</sup>C-Raclopride | 27 | Fig 4 | VST<br>Caudate nucleus<br>Putamen |  |  |
| Wang | 2001 | USA | <sup>11</sup>C-Raclopride | 10 | Fig 2 | Striatum |  |  |
| Steele | 2010 | USA | <sup>11</sup>C-Raclopride | 5 | Fig 4 | Striatum | Fig 4 | Striatum |
| Gaiser | 2016 | USA | <sup>11</sup>C-PHNO | 42 | Fig 2 | VST | Mean<br>BP<sub>ND</sub> | VST<br>Caudate nucleus<br>Putamen |
| Volkow | 2008 | USA | <sup>11</sup>C-Raclopride | 22 |  |  | Mean<br>BP<sub>ND</sub> | Striatum |
| Karlsson | 2016 | Finland | <sup>11</sup>C-Raclopride | 16 |  |  | Fig 2 | VST<br>Caudate nucleus<br>Putamen |
| Guo | 2014 | USA | <sup>18</sup>F-Fallypride | 43 |  |  | Mean<br>BP<sub>ND</sub> | NAc<br>Caudate nucleus<br>Putamen |
| Dunn | 2010 | USA | <sup>18</sup>F-Fallypride | 5 |  |  | Mean<br>BP<sub>ND</sub> | VST<br>Caudate nucleus<br>Putamen |

ROI, region of interest; VST, ventral striatum; BP<sub>ND</sub>, binding potential; NAc, nucleus accumbens

### Association Between BMI and BP_ND_

#### Method 1

Five studies involving 119 subjects were included in this analysis. ^11^C-raclopride was used in four studies, regardless of the ROIs included (Caravaggio et al., 2015; Karlsson et al., 2015; Steele et al., 2010; Wang et al., 2001) (Figure 2A). Fitting a LOWESS curve of BP_ND_ from VST was performed with data from two studies (Caravaggio et al., 2015; Karlsson et al., 2015). The curve of BP_ND_ of the VST went up and down at a BMI of 20–30 kg/m^2^ and became flat at a BMI of >40 kg/m^2^ (Figure 2B). Linear regression analysis was performed with the BP_ND_ of the striatum from two studies (n = 20) (Steele et al., 2010; Wang et al., 2001). The BP_ND_ of the striatum was negatively associated with BMI (p = 0.0158; *y* = −0.02013\**x* + 3.539) (Figure 2C). ^11^C-PHNO data were extracted from two studies (n = 68) (Caravaggio et al., 2015; Gaiser et al., 2016). The linear regression analysis showed a positive association of BMI with the BP_ND_ of VST (p < 0.0001; *y* = 0.06502\**x* + 2.387) (Figure 3).

**Figure 2.**
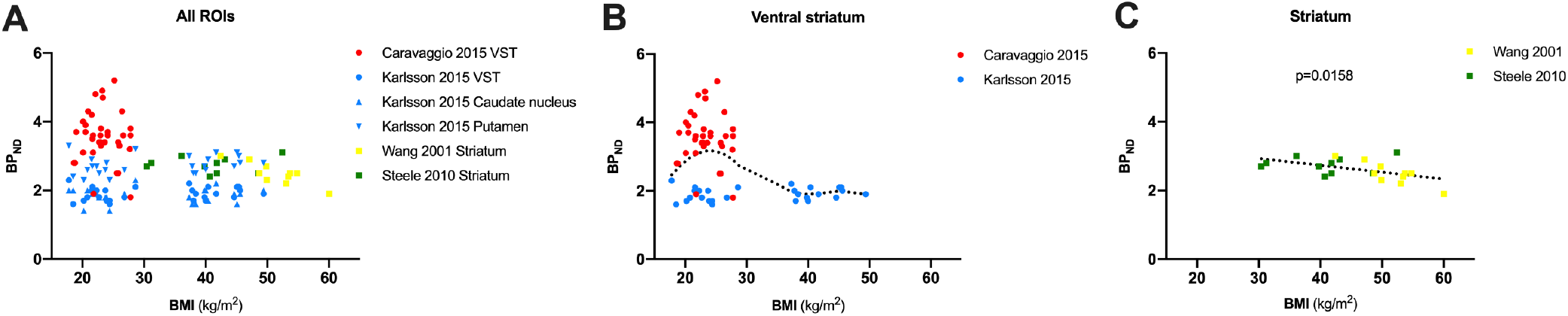
Scatter plots of the BP_ND_ of ^11^C-raclopride and BMI; A. All ROIs; B. Ventral striatum; C. Striatum.

**Figure 3.**
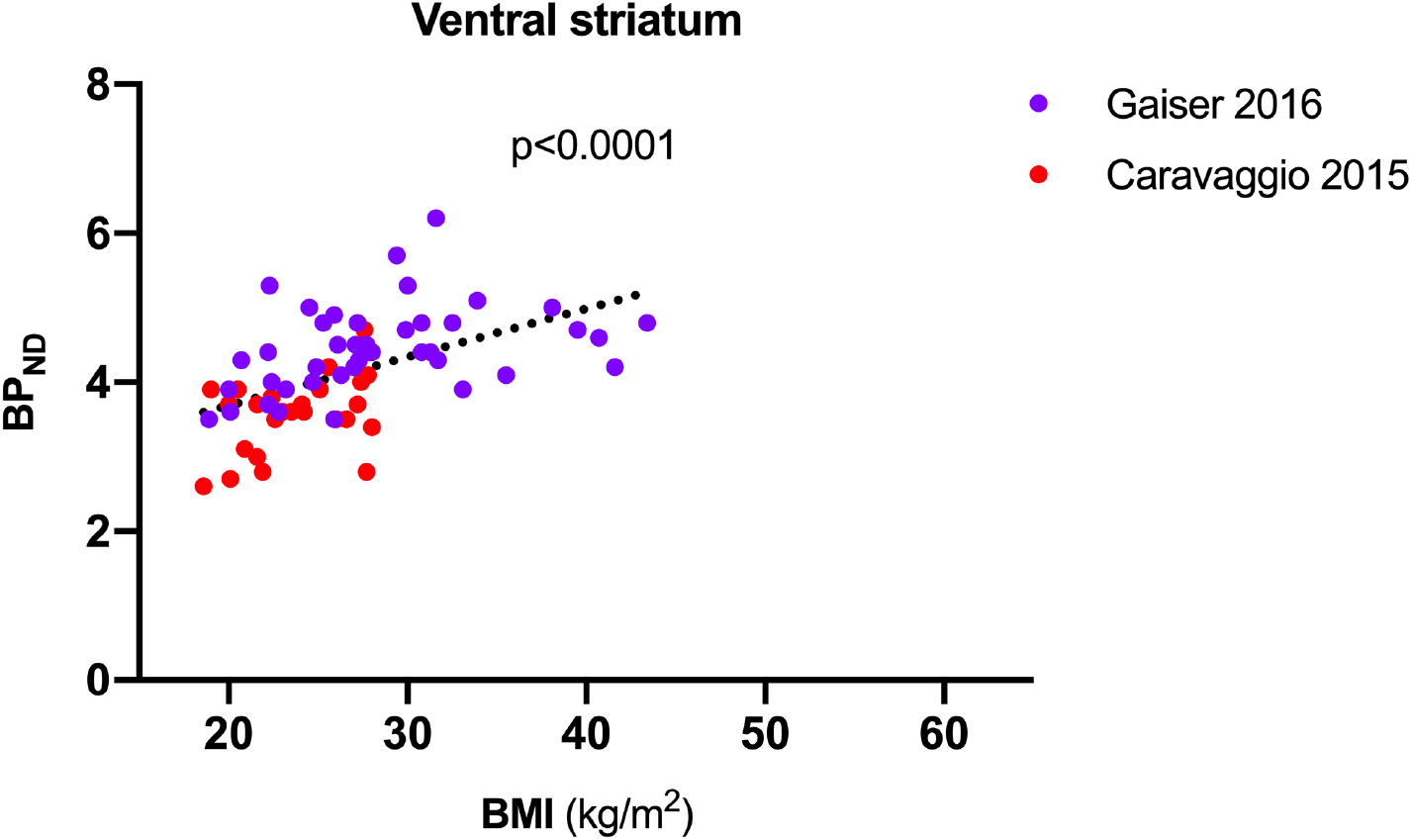
Scatter plots and regression line of the BP_ND_ of ^11^C-PHNO and BMI.

#### Method 2

Seven studies including 168 subjects were included in this analysis (Caravaggio et al., 2015; Dunn et al., 2010; Gaiser et al., 2016; Guo et al., 2014; Karlsson et al., 2016; Steele et al., 2010; Volkow et al., 2008). The mean BP_ND_ and BMI values were extracted directly from the results of four studies (Dunn et al., 2010; Gaiser et al., 2016; Guo et al., 2014; Volkow et al., 2008). The mean BMI and BP_ND_ values were calculated after extracting each BMI and corresponding BP_ND_ from figures in three studies (Caravaggio et al., 2015; Karlsson et al., 2016; Steele et al., 2010). ^11^C-raclopride was used in four studies (Caravaggio et al., 2015; Karlsson et al., 2016; Steele et al., 2010; Volkow et al., 2008), and each of ^11^C-PHNO (Caravaggio et al., 2015; Gaiser et al., 2016) and ^18^F-Fallypride (Dunn et al., 2010; Guo et al., 2014) was extracted from two studies (Figure 4). The distribution of the mean BMI and BP_ND_ values for ^11^C-raclopride showed a similar pattern to that of method 1. Scatter plots of ^11^C-PHNO and ^18^F-fallypride showed a higher mean BP_ND_ value with a higher mean BMI value. As there were less than five studies for each ROI and radiopharmaceutical, the regression analysis could not be performed.

**Figure 4.**
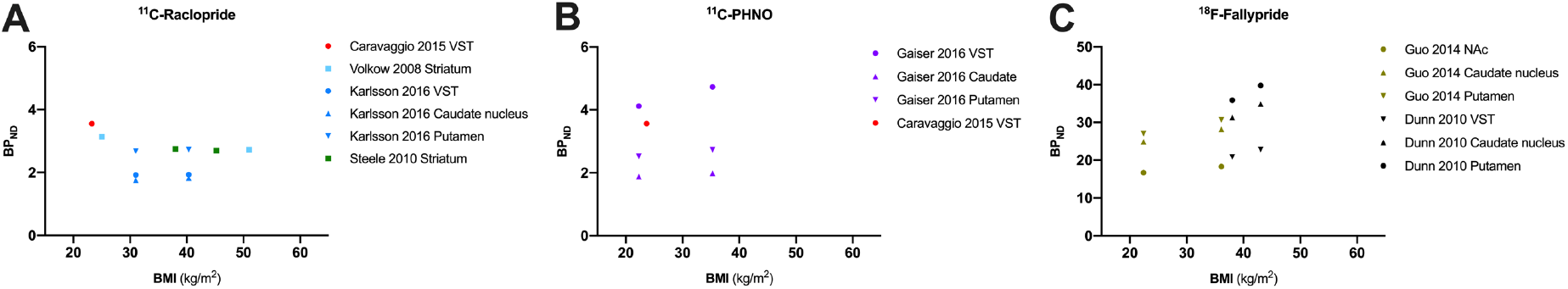
Scatter plots of the mean BP_ND_ and BMI; A. ^11^C-raclopride; B. ^11^C-PHNO; ^18^F-fallypride.

## DISCUSSION

Obesity is caused by an imbalance between the energy intake and expenditure (Bellisle, Drewnowski, Anderson, Westerterp-Plantenga, & Martin, 2012). The brain plays an important role in the primary center of this balance (Levin & Routh, 1996). The eating behavior is regulated mainly by the homeostatic (hypothalamus) and hedonic systems (striatum) of the brain; both the systems are closely linked to each other (van Galen et al., 2018). The hypothalamus plays a central role in maintaining the physiologic requirements of the body through regulatory neuropeptides (Khanh, Choi, Moh, Kinyua, & Kim, 2014; van Vliet-Ostaptchouk, Hofker, van der Schouw, Wijmenga, & Onland-Moret, 2009). The eating behavior is regulated by the reward system of the striatum (Khanh et al., 2014). Neurotransmitters, such as dopamine, opioid, and serotonin, mediate the hedonic functions of this reward system (Nummenmaa et al., 2018; van Galen et al., 2018). Among them, dopamine plays a main role in modulating motivation and processing rewards (Wang et al., 2011). There is no direct method to measure dopamine levels in the human brain. PET using DR radiopharmaceuticals has been used to understand the dopaminergic pathway in the brain. To elucidate the association between obesity and DRs, we adopted two methods in this meta-analysis.

In this study, BMI showed an inverse parabolic relationship with DR availability measured using ^11^C-raclopride from the VST in normal-weight or overweight subjects (BMI: <30 kg/m^2^), although neither of the two studies conclude any significant association of BMI with BP_ND_ (Caravaggio et al., 2015; Karlsson et al., 2015). DR availability increased till a BMI of approximately 25 kg/m^2^ in the mid-range between 20 and 30 kg/m^2^. However, among subjects with a BMI of approximately 25–30 kg/m^2^, those with a higher BMI score tended to have lower BP_ND_. There are two major hypotheses regarding the role of dopamine in obesity. The first hypothesis, dopamine hyperresponsiveness, explains that there is a hypersensitivity to rewards, and it is related to an increased behavioral salience toward food, resulting in the excessive intake of highly palatable foods (Kessler, Zald, Ansari, Li, & Cowan, 2014; Verbeken, Braet, Lammertyn, Goossens, & Moens, 2012). This might be consistent with our results of higher BP_ND_ with higher BMI in normal-weight subjects (BMI: <25 kg/m^2^). The second hypothesis, reward deficiency syndrome, states that subjects who are insensitive to rewards overeat to increase their endogenous dopamine levels to a normal amount of pleasure (Kessler et al., 2014; Verbeken et al., 2012), resulting in decreased DR availability, which is consistent with the results for overweight subjects (25 kg/m^2^ ≤ BMI < 30 kg/m^2^) in this study. As the subjects with a BMI of 29–37 kg/m^2^ were not included in two studies (Caravaggio et al., 2015; Karlsson et al., 2015), DR availabilities of those subjects with obesity class I (30 kg/m^2^ ≤ BMI < 35 kg/m^2^) could not be assessed in this meta-analysis. In subjects with obesity class II or III, the curve of BP_ND_ became flat, regardless of the BMI. However, in the studied by Wang et al. (Wang et al., 2001), and Steele et al. (Steele et al., 2010), DR availability of the striatum was negatively associated with a BMI of 30–60 kg/m^2^. This inconsistency could be derived from either the difference in ROIs between the VST and the striatum or distribution of BMI data. The BMI in the study by Wang et al. (Wang et al., 2001) and Steele et al. (Steele et al., 2010) ranged widely from 30 to 60 kg/m^2^, with 30% (6 of 20) subjects having a BMI of >50 kg/m^2^, while the BMI in the study by Karlsson et al. (Karlsson et al., 2015) ranged between 37 and 50 kg/m^2^, with subjects with less severe obesity included.

The DR availability measured using BP_ND_ of ^11^C-PHNO was strongly associated with BMI. In the studies by Gaiser et al. (Gaiser et al., 2016) and Caravaggio et al. (Caravaggio et al., 2015), subjects with a higher BMI had a higher DR availability measured using ^11^C-PHNO. These differences might result from the characteristics of ^11^C-raclopride and ^11^C-PHNO. Both ^11^C-raclopride and ^11^C-PHNO are sensitive to changes in endogenous dopamine levels (Shotbolt et al., 2012), with the latter being more sensitive than the former (Gallezot et al., 2014). In addition, ^11^C-PHNO, a D2/3R agonist, has greater binding in D3R, while ^11^C-raclopride, a D2/3R antagonist, predominantly estimates D2R (Graff-Guerrero et al., 2008). Therefore, the results of DR availability measured using both ^11^C-raclopride and ^11^C-PHNO from the mixed D2/3R area of the VST might be inconsistent (Gaiser et al., 2016). This inconsistency also suggests that D2R and D3R may play different roles in the reward system with regard to obesity (Gaiser et al., 2016).

In a previous study, van Galen et al. compared the DR availability of obsess subjects and lean subjects, regardless of radiopharmaceuticals, including ^11^C-PHNO, ^18^F-fallypride, ^11^C-raclopride, ^11^C-NMB, and ^123^I-IBZM (van Galen et al., 2018). There was a non-linear relationship between BMI and DR availability; thee DR availability increased in moderate obesity (BMI: approximately 35 kg/m^2^) and the DR availability decreased with the progression of obesity (BMI >40 kg/m^2^) (van Galen et al., 2018). Although with the mean BMI and BP_ND_ values may not reflect the association between obesity and DR availability in each subject, we presented the plots to investigate the trend between them according to the radiopharmaceuticals used in method 2. Different from the study by van Galen et al. (van Galen et al., 2018), DR availability goes down between BMI scores of 20 and 30 kg/m^2^ in both methods 1 and 2 in this study, which might reflect the downregulation of DR from overweight BMI before the onset of obesity. In addition, subjects with a higher BMI seemed to have a higher DR availability measured using both ^11^C-PHNO and ^18^F-fallypride, similar to the result from method 1. As each radiopharmaceutical might represent different aspects of DR in the brain, these results should be interpreted separately.

This study has several limitations. First, a small number of studies was included in this meta-analysis. Therefore, each of the two studies could be included in the analysis of method 1 according to ROIs. Second, radiopharmaceuticals, image processing, and data analysis techniques are different across the studies included in this meta-analysis, which might impact BP_ND_. As each of the two studies were included in each analysis, the heterogeneity of the study design could not be solved in this meta-analysis. Third, although two authors extracted the BMI and corresponding BP_ND_ values from the figures in the studies, the data may be inaccurate.

## CONCLUSION

Dopamine plays a main role in the reward system with regard to obesity. Compared to normal-weight subjects, overweight and obese subjects had decreased DR availability. However, this tendency should be interpreted carefully with regard to the radiopharmaceuticals and ROIs.

## ACKNOWLEDGEMENT

None

## DATA AVAILABILITY

The datasets generated during the current study are available from the corresponding author on reasonable request.

## REFERENCES

Alzahrani, B., Iseli, T. J., & Hebbard, L. W. (2014). Non-viral causes of liver cancer: does obesity led inflammation play a role? Cancer Lett, 345(2), 223–229. doi:10.1016/j.canlet.2013.08.036

Baik, J. H. (2013). Dopamine signaling in reward-related behaviors. Front Neural Circuits, 7, 152. doi:10.3389/fncir.2013.00152

Bellisle, F., Drewnowski, A., Anderson, G. H., Westerterp-Plantenga, M., & Martin, C. K. (2012). Sweetness, satiation, and satiety. J Nutr, 142(6), 1149S–1154S. doi:10.3945/jn.111.149583

Berridge, K. C. (1996). Food reward: brain substrates of wanting and liking. Neurosci Biobehav Rev, 20(1), 1–25.

Burke, G. L., Bertoni, A. G., Shea, S., Tracy, R., Watson, K. E., Blumenthal, R. S., … Carnethon, M. R. (2008). The impact of obesity on cardiovascular disease risk factors and subclinical vascular disease: the Multi-Ethnic Study of Atherosclerosis. Arch Intern Med, 168(9), 928–935. doi:10.1001/archinte.168.9.928

Caravaggio, F., Raitsin, S., Gerretsen, P., Nakajima, S., Wilson, A., & Graff-Guerrero, A. (2015). Ventral striatum binding of a dopamine D2/3 receptor agonist but not antagonist predicts normal body mass index. Biol Psychiatry, 77(2), 196–202. doi:10.1016/j.biopsych.2013.02.017

Dunn, J. P., Cowan, R. L., Volkow, N. D., Feurer, I. D., Li, R., Williams, D. B., … Abumrad, N. N. (2010). Decreased dopamine type 2 receptor availability after bariatric surgery: preliminary findings. Brain Res, 1350, 123–130. doi:10.1016/j.brainres.2010.03.064

Eisenstein, S. A., Antenor-Dorsey, J. A., Gredysa, D. M., Koller, J. M., Bihun, E. C., Ranck, S. A., … Hershey, T. (2013). A comparison of D2 receptor specific binding in obese and normal-weight individuals using PET with (N-[(11)C]methyl)benperidol. Synapse, 67(11), 748–756. doi:10.1002/syn.21680

Gaiser, E. C., Gallezot, J. D., Worhunsky, P. D., Jastreboff, A. M., Pittman, B., Kantrovitz, L., … Matuskey, D. (2016). Elevated Dopamine D2/3 Receptor Availability in Obese Individuals: A PET Imaging Study with [(11)C](+)PHNO. Neuropsychopharmacology, 41(13), 3042–3050. doi:10.1038/npp.2016.115

Gallezot, J. D., Kloczynski, T., Weinzimmer, D., Labaree, D., Zheng, M. Q., Lim, K., … Cosgrove, K. P. (2014). Imaging nicotine- and amphetamine-induced dopamine release in rhesus monkeys with [(11)C]PHNO vs [(11)C]raclopride PET. Neuropsychopharmacology, 39(4), 866–874. doi:10.1038/npp.2013.286

Graff-Guerrero, A., Willeit, M., Ginovart, N., Mamo, D., Mizrahi, R., Rusjan, P., … Kapur, S. (2008). Brain region binding of the D2/3 agonist [11C]-(+)-PHNO and the D2/3 antagonist [11C]raclopride in healthy humans. Hum Brain Mapp, 29(4), 400–410. doi:10.1002/hbm.20392

Gu, W., Chen, C., & Zhao, K. N. (2013). Obesity-associated endometrial and cervical cancers. Front Biosci (Elite Ed), 5, 109–118.

Gukovsky, I., Li, N., Todoric, J., Gukovskaya, A., & Karin, M. (2013). Inflammation, autophagy, and obesity: common features in the pathogenesis of pancreatitis and pancreatic cancer. Gastroenterology, 144(6), 1199–1209 e1194. doi:10.1053/j.gastro.2013.02.007

Guo, J., Simmons, W. K., Herscovitch, P., Martin, A., & Hall, K. D. (2014). Striatal dopamine D2-like receptor correlation patterns with human obesity and opportunistic eating behavior. Mol Psychiatry, 19(10), 1078–1084. doi:10.1038/mp.2014.102

Heimer, L. (1978). Limbic Mechanisms. New York: Plenum Press.

Karlsson, H. K., Tuominen, L., Tuulari, J. J., Hirvonen, J., Parkkola, R., Helin, S., … Nummenmaa, L. (2015). Obesity is associated with decreased mu-opioid but unaltered dopamine D2 receptor availability in the brain. J Neurosci, 35(9), 3959–3965. doi:10.1523/JNEUROSCI.4744-14.2015

Karlsson, H. K., Tuulari, J. J., Tuominen, L., Hirvonen, J., Honka, H., Parkkola, R., … Nummenmaa, L. (2016). Weight loss after bariatric surgery normalizes brain opioid receptors in morbid obesity. Mol Psychiatry, 21(8), 1057–1062. doi:10.1038/mp.2015.153

Kenny, P. J. (2011). Common cellular and molecular mechanisms in obesity and drug addiction. Nat Rev Neurosci, 12(11), 638–651. doi:10.1038/nrn3105

Kessler, R. M., Zald, D. H., Ansari, M. S., Li, R., & Cowan, R. L. (2014). Changes in dopamine release and dopamine D2/3 receptor levels with the development of mild obesity. Synapse, 68(7), 317–320. doi:10.1002/syn.21738

Khanh, D. V., Choi, Y. H., Moh, S. H., Kinyua, A. W., & Kim, K. W. (2014). Leptin and insulin signaling in dopaminergic neurons: relationship between energy balance and reward system. Front Psychol, 5, 846. doi:10.3389/fpsyg.2014.00846

Lam, D. D., Garfield, A. S., Marston, O. J., Shaw, J., & Heisler, L. K. (2010). Brain serotonin system in the coordination of food intake and body weight. Pharmacol Biochem Behav, 97(1), 84–91. doi:10.1016/j.pbb.2010.09.003

Levin, B. E., & Routh, V. H. (1996). Role of the brain in energy balance and obesity. Am J Physiol, 271(3 Pt 2), R491–500. doi:10.1152/ajpregu.1996.271.3.R491

Mijovic, T., How, J., & Payne, R. J. (2011). Obesity and thyroid cancer. Front Biosci (Schol Ed), 3, 555–564.

Morton, G. J., Meek, T. H., & Schwartz, M. W. (2014). Neurobiology of food intake in health and disease. Nat Rev Neurosci, 15(6), 367–378. doi:10.1038/nrn3745

Na, S. Y., & Myung, S. J. (2012). [Obesity and colorectal cancer]. Korean J Gastroenterol, 59(1), 16–26.

Nummenmaa, L., Saanijoki, T., Tuominen, L., Hirvonen, J., Tuulari, J. J., Nuutila, P., & Kalliokoski, K. (2018). mu-opioid receptor system mediates reward processing in humans. Nat Commun, 9(1), 1500. doi:10.1038/s41467-018-03848-y

Organization, W. H. (2018). Obesity and overweight. Retrieved from http://www.who.int/news-room/fact-sheets/detail/obesity-and-overweight

Pak, K., Kim, S. J., & Kim, I. J. (2018). Obesity and Brain Positron Emission Tomography. Nucl Med Mol Imaging, 52(1), 16–23. doi:10.1007/s13139-017-0483-8

Pecina, S., Smith, K. S., & Berridge, K. C. (2006). Hedonic hot spots in the brain. Neuroscientist, 12(6), 500–511. doi:10.1177/1073858406293154

Ravussin, E., & Bogardus, C. (2000). Energy balance and weight regulation: genetics versus environment. Br J Nutr, 83 Suppl 1, S17–20.

Shotbolt, P., Tziortzi, A. C., Searle, G. E., Colasanti, A., van der Aart, J., Abanades, S., … Rabiner, E. A. (2012). Within-subject comparison of [(11)C]-(+)-PHNO and [(11)C]raclopride sensitivity to acute amphetamine challenge in healthy humans. J Cereb Blood Flow Metab, 32(1), 127–136. doi:10.1038/jcbfm.2011.115

Steele, K. E., Prokopowicz, G. P., Schweitzer, M. A., Magunsuon, T. H., Lidor, A. O., Kuwabawa, H., … Wong, D. F. (2010). Alterations of central dopamine receptors before and after gastric bypass surgery. Obes Surg, 20(3), 369–374. doi:10.1007/s11695-009-0015-4

Val-Laillet, D., Aarts, E., Weber, B., Ferrari, M., Quaresima, V., Stoeckel, L. E., … Stice, E. (2015). Neuroimaging and neuromodulation approaches to study eating behavior and prevent and treat eating disorders and obesity. Neuroimage Clin, 8, 1–31. doi:10.1016/j.nicl.2015.03.016

van Galen, K. A., Ter Horst, K. W., Booij, J., la Fleur, S. E., & Serlie, M. J. (2018). The role of central dopamine and serotonin in human obesity: lessons learned from molecular neuroimaging studies. Metabolism, 85, 325–339. doi:10.1016/j.metabol.2017.09.007

van Vliet-Ostaptchouk, J. V., Hofker, M. H., van der Schouw, Y. T., Wijmenga, C., & Onland-Moret, N. C. (2009). Genetic variation in the hypothalamic pathways and its role on obesity. Obes Rev, 10(6), 593–609. doi:10.1111/j.1467-789X.2009.00597.x

Verbeken, S., Braet, C., Lammertyn, J., Goossens, L., & Moens, E. (2012). How is reward sensitivity related to bodyweight in children? Appetite, 58(2), 478–483. doi:10.1016/j.appet.2011.11.018

Volkow, N. D., Wang, G. J., Telang, F., Fowler, J. S., Thanos, P. K., Logan, J., … Pradhan, K. (2008). Low dopamine striatal D2 receptors are associated with prefrontal metabolism in obese subjects: possible contributing factors. Neuroimage, 42(4), 1537–1543. doi:10.1016/j.neuroimage.2008.06.002

Volkow, N. D., Wang, G. J., Tomasi, D., & Baler, R. D. (2013). The addictive dimensionality of obesity. Biol Psychiatry, 73(9), 811–818. doi:10.1016/j.biopsych.2012.12.020

Wang, G. J., Geliebter, A., Volkow, N. D., Telang, F. W., Logan, J., Jayne, M. C., Fowler, J. S. (2011). Enhanced striatal dopamine release during food stimulation in binge eating disorder. Obesity (Silver Spring), 19(8), 1601–1608. doi:10.1038/oby.2011.27

Wang, G. J., Volkow, N. D., Logan, J., Pappas, N. R., Wong, C. T., Zhu, W., Fowler, J. S. (2001). Brain dopamine and obesity. Lancet, 357(9253), 354–357.

